# Single-cell transcriptomics reveals conserved embryonic-like cell populations in different human cancers

**DOI:** 10.64898/2025.12.11.693590

**Authors:** Anna A. Khozyainova, Vera G. Subrakova, Maxim E. Menyailo, Daria I. Zhigalina, Tatiana N. Kireeva, Adelya A. Galiakberova, Mikhail A. Berestovoy, Elena E. Kopantseva, Anastasia A. Korobeynikova, Erdem B. Dashinimaev, Dmitry M. Loos, Maria S. Tretyakova, Ustinia A. Bokova, Nikolay A. Skryabin, Evgeny V. Denisov

## Abstract

**Background:** Embryonic development is governed by coordinated transcriptional programs that shape cellular differentiation and can be reactivated in cancer. However, whether tumors of distinct ontogenetic origins contain conserved tumor cell populations that recapitulate features of these developmental programs remains unclear.

**Methods:** Here, we integrated single-cell RNA sequencing data from embryoid bodies (EBs), an in vitro model of early human development, and three ontogenetically distinct cancers (lung adenocarcinoma, melanoma, and medulloblastoma) to identify conserved tumor cell populations, whose features were further validated in an independent pan-cancer single-cell RNA sequencing atlas and by spatial transcriptomics.

**Results:** Across three cancer types, integrative analysis identified conserved tumor cell populations transcriptionally similar to EB-derived cells. The populations were marked by pronounced proliferation, recurrent expression of *UBE2C*, *TOP2A*, *BIRC5*, and *NUSAP1*, coordinated E2F-TFDP activity, and enrichment of cell-cycle, DNA repair, and mitotic programs together with low differentiation, high inferred developmental potential, and early pseudotime positioning. A proliferation-associated signature derived from these populations recurred across multiple cancer types and was more frequently detected in tumor than in normal cells, demonstrating altered expression of several cancer-related genes. Spatial transcriptomics localized the conserved tumor cell populations and their signature to premalignant bronchial lesions and discrete regions of lung adenocarcinoma tissue.

**Conclusions:** Together, our findings identify conserved tumor cell populations across cancers of distinct origins with transcriptional similarities to EB-derived cells including pronounced proliferation, low differentiation, and high inferred developmental potential and provide a foundation for developing new diagnostic and therapeutic solutions.

Clinical trial number: not applicable.

**Graphical abstract:** 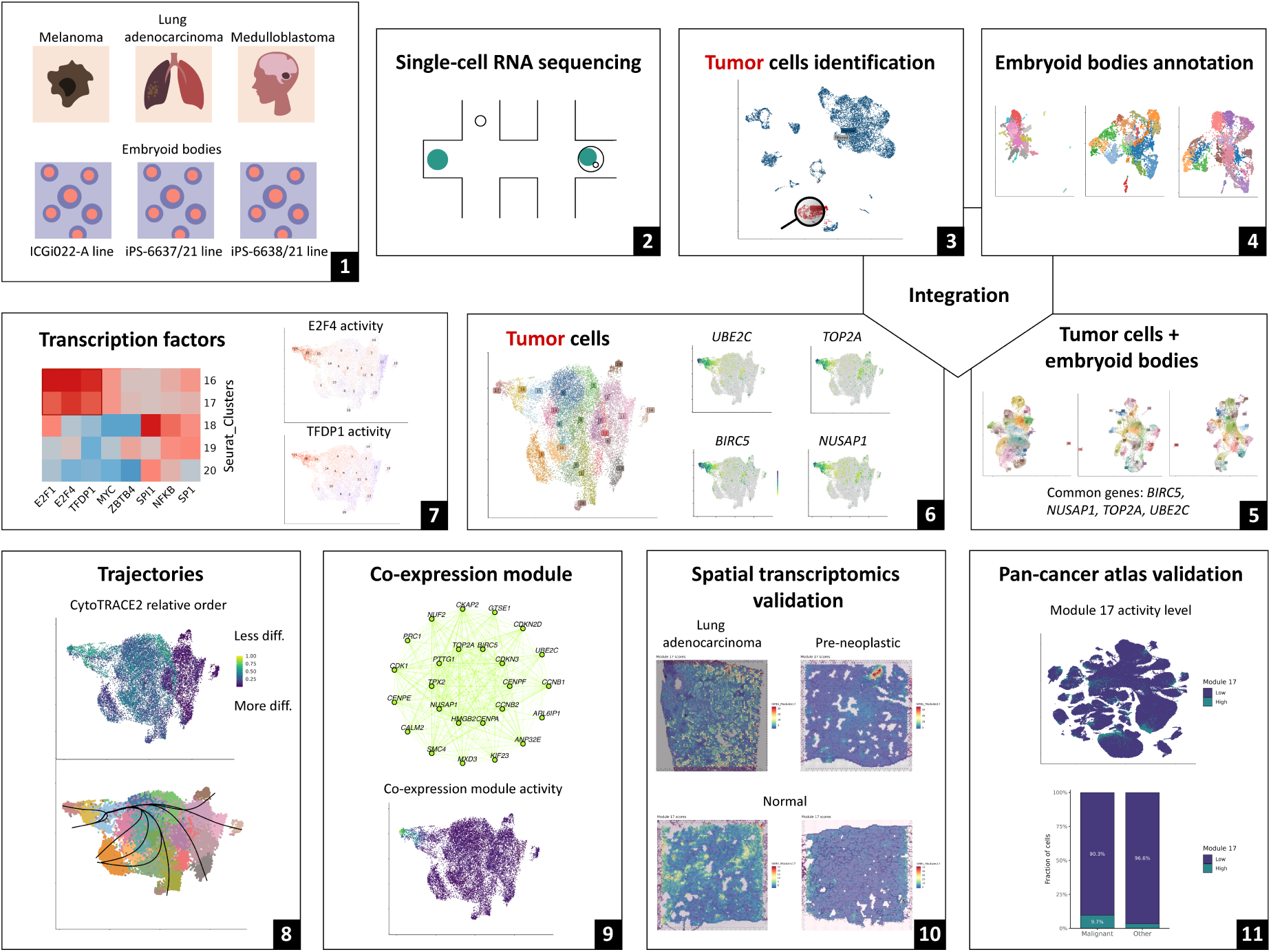

## Background

Embryonic development is governed by coordinated transcriptional programs that direct lineage specification, morphogenesis, and organogenesis through dynamic transitions between cellular states (1, 2). These programs are established through the remodeling of gene regulatory and epigenetic circuits and are regulated by networks of signaling cascades and transcription factors (3, 4). Contemporary single-cell and spatial transcriptomics approaches have enabled the characterization of universal transcriptional states and key regulatory axes in early human development (5, 6).

Remarkably, many of the molecular circuits that govern embryonic development are reactivated in pathological contexts, most notably in cancer. Increasing evidence suggests that oncogenesis co-opts developmental programs to fuel uncontrolled proliferation, invasion, metastasis, and therapy resistance (7, 8). This convergence highlights a profound biological parallel: the mechanisms that orchestrate normal development could be aberrantly deployed to drive malignant progression, including angiogenesis, epithelial-mesenchymal transition (EMT), acquisition of stem-like properties, and immune evasion, among other processes reminiscent of early embryogenesis (9–12).

Recently, single-cell transcriptomic studies across diverse cancer types have shown that tumor cells adopt states resembling those in normal development and remain restricted to the lineage hierarchies of their tissues of origin (13–16). This perspective has given rise to the concept of developmental constraints, in which cancer cells, despite their genomic instability and phenotypic flexibility, remain restricted to transcriptional states accessible within the organism’s original developmental map (17). Tumor cells do not generate arbitrary cell identities; rather, they traverse adjacent progenitor and differentiated states that reflect lineage relationships established during embryogenesis. Viewing tumor cell states through the lens of embryonic developmental programs could offer new insights, for example, explaining why cancers typically follow lineage-dependent trajectories imposed by their tissue of origin. More importantly, this approach can reveal unique biological pathways tumor cells depend on, pointing us toward new strategies to stop tumor growth and overcome treatment resistance (11, 18, 19). However, human post-implantation embryogenesis remains poorly understood because direct observation and experimental manipulation at this stage are severely limited by ethical and technical constraints, which represents a critical barrier to decoding and targeting cancer-specific vulnerabilities (20, 21).

One potential solution involves the use of embryoid bodies (EBs), which recapitulate key features of early development, including the presence of cells from the three germ layers and the expression of characteristic markers (22). In particular, studies of cell mechanics in EBs demonstrate that upon aggregation, pluripotent cells self-organize into structured, multicellular assemblies that undergo morphogenetic processes reminiscent of early embryo development, including cellular differentiation, EMT, lumen formation, and formation of primitive embryonic-like tissues (23). Moreover, recent data indicate that human EBs can generate a diverse spectrum of cell types and developmental trajectories, and reproduce aspects of lineage specification and cell-type heterogeneity observed *in vivo* (24, 25). Thus, the EBs system offers a tracable, ethically acceptable, and experimentally controllable platform to model events of early human embryogenesis, bridging the gap between in vivo embryo research and in vitro cellular models. Consequently, EBs could serve as a developmental reference framework enabling direct comparison of tumor transcriptional states with those characteristic of early embryonic cell populations.

The present study aims to identify conserved tumor cell populations, transcriptional programs, and regulatory axes associated with embryonic development across cancers of distinct origins. To achieve this, we compared the transcriptional profiles of embryoid bodies and three ontogenetically distinct tumors (lung adenocarcinoma, melanoma, and medulloblastoma) using integrative single-cell RNA sequencing (scRNA-seq) analysis, followed by validation in an independent pan-cancer scRNA-seq atlas and by spatial transcriptomics.

## Methods

### Embryoid bodies preparation

#### EBs from ICGi022-A cell line

EBs were obtained from induced pluripotent stem cells (iPSCs) of a healthy individual (ICGi022-A cell line), provided by the Institute of Cytology and Genetics of the Siberian Branch of the Russian Academy of Sciences, Novosibirsk (26). iPSC colonies of the ICGi022-A cell line were detached using 0.15% type IV collagenase (Thermo Fisher Scientific, USA) and transferred to 30-mm dishes coated with 1% agarose to initiate undirected differentiation. These iPSCs were cultivated for 18 days in iPSC medium without basic fibroblast growth factor (bFGF): Dulbecco’s Modified Eagle Medium (DMEM)/Nutrient Mixture F-12 (Thermo Fisher Scientific, USA) supplemented with 15% Knockout Serum Replacement (KoSR), 1× MEM Non-Essential Amino Acids Solution (Gibco, USA), 0.1 mM 2-mercaptoethanol (Gibco, USA), 1× penicillin-streptomycin (Gibco, USA), and 1× GlutaMAX.

#### EBs from iPS-6638/21 and iPS-6637/21 cell lines

iPS-6638/21 and iPS-6637/21 cell lines were derived from healthy female and male donor fibroblasts using a Sendai virus reprogramming kit. iPSCs were seeded at a density of 2000 cells/cm² on 6 cm Petri dishes coated with 2% Matrigel in mTeSR Plus medium supplemented with 5 µM ROCK Inhibitor Y-27632 (STEMCELL Technologies, Canada). Once the iPSCs formed individual small colonies, the dishes were incubated with a dispase solution (3 mg/ml in DMEM/F12; HiMedia, India) at 37°C for 20-30 minutes to detach the colonies as intact aggregates. The detached colonies were transferred to a 15 ml tube with DPBS without calcium and magnesium and centrifuged at 100 g for 1-2 min. The pellet was carefully resuspended in mTeSR Plus and transferred to ultra-low binding Petri dishes for suspension culture in mTeSR Plus supplemented with 5 µM ROCK Inhibitor Y-27632 (STEMCELL Technologies, Canada). A partial medium change was performed every other day, gradually replacing mTeSR Plus with embryoid body medium including DMEM/F12 (PanEco, Russia), 20% fetal bovine serum (FBS) (Capricorn Scientific, Germany), 2 mM Glutargo (Capricorn Scientific, Germany), 1 mM Sodium Pyruvate (Gibco, USA), 0.1 mM β-Mercaptoethanol (Sigma-Aldrich, USA), 1× NEAA (Capricorn Scientific, Germany), and PenStrep (50 U/ml; 50 µg/ml; Gibco, USA).

### Immunofluorescence staining of EBs

EBs of the ICGi022-A cell line were transferred to gelatin-coated 6-well plates with mouse embryonic fibroblasts for an additional eight days. EBs were fixed in 3% paraformaldehyde and incubated with primary antibodies overnight at 4 °C. Following washes with PBS containing 0.1% Tween-20, samples were incubated with Alexa Fluor-conjugated secondary antibodies diluted in blocking buffer. Nuclei were counterstained with DAPI (Sigma-Aldrich, USA) in antifade mounting medium (Vectashield, Vector Laboratories, USA). Fluorescence images were acquired using an Axio Imager Z2 microscope (Zeiss, Germany) equipped with ISIS software (MetaSystems, Germany). After 14 days of suspension culture, EBs from the iPS-6638/21 and iPS-6637/21 cell lines were collected and plated into wells of a 24-well plate, pre-coated with 2% Matrigel, at a density of 6 EBs per well in the embryoid body medium. The medium was changed 2-3 times per week. Following additional 14 days of spontaneous differentiation, the resulting cultures were fixed for subsequent immunocytochemical analysis as described above. The antibodies used for immunocytochemistry are listed in Supplementary Table 1.

### Generation of single-cell suspensions

#### EBs from ICGi022-A cell line

The EBs were transported on ice in 2 mL of DMEM/F12 culture medium (Thermo Fisher Scientific, USA) supplemented with 20% FBS (Global Kang Biotechnology, China). The experimental sample consisted of approximately 20-25 EBs. The EBs were centrifuged at 300g for 10 min, and the supernatant was removed and replaced with 5 mL of DMEM/F12 containing the enzyme cocktail from the Human Tumor Dissociation Kit (100 µL Enzyme H, 50 µL Enzyme R, and 12.5 µL Enzyme A; Miltenyi Biotec, Germany). The EBs were incubated for 30 min at 37 °C in the gentleMACS Octo Dissociator with Heaters (Miltenyi Biotec, Germany). After dissociation, the cell suspension was passed through a 70-µm cell strainer, centrifuged at 300g for 5 min, and resuspended in 1 mL of fresh DMEM/F12. Cells were counted using 0.4% Trypan Blue Solution (Thermo Fisher Scientific, USA) on a LUNA-II Automated Cell Counter (Logos Biosystems, South Korea). The experimental sample yielded approximately 1 × 10⁶ cells, with a viability of ∼80%. The scRNA-seq experiment included 16500 cells.

#### EBs from iPS-6638/21 and iPS-6637/21 cell lines

The EBs were transported on ice in 2 mL of the embryoid body medium. Each experimental sample contained approximately 20-25 EBs. The EBs were centrifuged at 300g for 5 minutes, and their culture medium was replaced with 2 mL of DMEM High Glucose (ServiceBio, China) with the enzyme cocktail from the Human Tumor Dissociation Kit (100 µL Enzyme H, 50 µL Enzyme R, 12.5 µL Enzyme A; Miltenyi Biotec, Germany). The EBs were incubated for 30 minutes at 37 °C and 1400 rpm in a TS-100C Smart thermoshaker (Biosan, Latvia). After the incubation, the resulting dissociated cells were passed through a 40-µm cell strainer, centrifuged at 300g for 5 minutes, and resuspended in 1 mL of fresh DMEM High Glucose (ServiceBio, China). The cells were counted using an acridine orange/propidium iodide stain buffer (Logos Biosystems, South Korea) on LUNA-FL Dual Fluorescence Cell Counter (Logos Biosystems, South Korea). The two experimental samples had around 10^6^ cells with approximately 71% viability. The scRNA-seq experiment included 16500 cells.

### Single-cell RNA sequencing library preparation

The single-cell transcriptome libraries were prepared using the Single Cell 3′ Reagent Kit v3 (10x Genomics, USA) according to the manufacturer’s protocol. GEM generation was performed on a Chromium iX/X instrument (10x Genomics, USA) for the EBs samples from the iPS-6638/21 and iPS-6637/21 lines and on a Chromium Controller (10x Genomics, USA) for the ICGi022-A EBs sample. Reverse transcription and cDNA amplification were carried out using a QuantGene 9600 thermal cycler (Bioer, China) for the EBs samples from the iPS-6638/21 and iPS-6637/21 lines, and a SimpliAmp Thermal Cycler (Thermo Fisher Scientific, USA) for the EBs sample from ICGi022-A line. Final libraries were diluted to 3.6 pM and sequenced on the GenolabM platform (GeneMind, China) with the following run configuration: Read 1 - 28 cycles, Read 2 - 90 cycles.

### Publicly available scRNA-seq datasets

Raw sequencing data used in this study were obtained from the Sequence Read Archive (SRA) (26). scRNA-seq data for lung adenocarcinoma were retrieved from the SRA Study SRP347260 and included samples SRR17008551, SRR17008552, and SRR17008559. Medulloblastoma scRNA-seq data were obtained from the SRA Study SRP274357, comprising samples SRR12354570, SRR12354583, and SRR12354584. Melanoma scRNA-seq data were retrieved from the SRA Study SRP389673, including samples SRR20791201, SRR20791205, and SRR20791206. A detailed overview of the tumor samples included in the analysis is shown in Supplementary Table 2.

### Single-cell RNA-seq data processing

Raw sequencing reads from lung adenocarcinoma, melanoma, medulloblastoma and EBs samples were analyzed with the Cell Ranger 7.1.0 pipeline (10x Genomics, USA). The resulting feature-barcode matrices were imported into the R programming environment (4.5.1) for further analysis with Seurat (5.3.0) (27). Doublet correction was performed using the scDblFinder tool (1.22.0) with default parameters (28). Cells with fewer than 500 detected genes or more than 20% mitochondrial content were excluded from the EBs datasets. For the tumor samples, cells with fewer than 200 detected genes, identified as doublets/multiplets, or with >20% mitochondrial content were excluded. Cell-level normalization and scaling were performed using SCTransform v2. The number of principal components (PCs) retained for downstream analysis was determined as follows: 50 PCs for EBs samples and 15 PCs for tumor samples. Neighbor graphs were constructed, and clustering was performed using the FindNeighbors and FindClusters functions with default parameters. Clustering resolution was set to 0.8 for tumor samples and to 1.2 for EBs samples. Dimensionality reduction and two-dimensional visualization were performed using uniform manifold approximation and projection (UMAP) on the selected PCs.

### Cell type annotation

The resulting clusters of tumor samples were annotated into major cell types with CellTypist (1.6.3) using the Human Lung Cell Atlas as a reference model for lung adenocarcinoma samples, the Adult Human Skin Atlas for melanoma samples, and the Adult Human Middle Temporal Gyrus Atlas for the medulloblastoma samples (29–32). In all samples, tumor cells were identified with SCEVAN (1.0.3) with immune cells used as a reference (33). Additionally, tumor cell annotation was confirmed with the inferCNV package (1.24.0) (34). Immune cells in each dataset (CD8 T cells, B cells, CD4 T cells, DC2, NK cells for lung adenocarcinoma and melanoma samples, Micro-PVM_2 for medulloblastoma samples) were designated as normal controls to establish a baseline CNV profile. Suggested tumor cells were analyzed with default parameters with cutoff set at 0.1.

Manual cell type annotation for EBs was performed based on differentially expressed genes (DEGs). Gene sets characteristic of each cluster were queried against the following databases and electronic resources: GeneAnalytics (35), GeneCards (36), LifeMap Discovery (37), WebGestalt (38), and The Human Protein Atlas (39, 40). Additionally, EBs samples were annotated with CellTypist using the Pan Fetal Human Atlas (41).

### Integration of EBs and cancer datasets

To investigate transcriptional similarities between early embryogenesis and oncogenesis, EBs cells and tumor cells from three cancer types, previously identified using SCEVAN, were integrated using the IntegrateLayers function with the RPCAIntegration method. Integrations were performed in two modes: pairwise integrations of tumor cells from each individual cancer type (lung adenocarcinoma, melanoma, and medulloblastoma separately) with EBs cells and a joint integration of tumor cells from all three cancer types. As a control analysis to exclude potential effects of cell cycle-associated variation, an additional joint integration of tumor cells from all three cancer types was performed with cell cycle effects regressed out. Following each integration, the data were scaled, and principal component analysis (PCA) was performed. Neighbor graphs were constructed using the first 30 PCs, followed by UMAP dimensionality reduction and Louvain clustering at resolution 1.0. UMAP feature plots showing gene expression were visualized with the scCustomize package (3.2.0) (42).

### Differential gene expression analysis

Identification of DEGs between clusters containing both tumor and EBs cells was performed using the FindConservedMarkers function with grouping.var set to a unique sample ID. For DEGs from clusters 16 and 17 from the integrated dataset of three cancers, FindConservedMarkers function with grouping.var set to a unique sample ID was similarly used. DEGs were identified using the following criteria: log2FC > 0.3 and adjusted p-value < 0.05 and were subsequentrly used for the Gene Ontology (GO) and Kyoto Encyclopedia of Genes and Genomes (KEGG) enrichment analyses. In addition, a list of DEGs from these clusters was identified with FindMarkers with default parameters, filtered to include DEGs with adjusted p-value < 0.05, and used for over-representation analysis of Molecular Signatures Database (MSigDB) Hallmark (H) gene sets analysis.

### Gene set enrichment analysis

Gene Set Enrichment Analysis (GSEA) was performed on the list of DEGs from clusters 16 and 17, as well as on the 25 hub genes from Module 17 using the clusterProfiler (4.16.0) package (43). The enricher function was used to test significant over-representation of MSigDB Hallmark (H) gene sets among significantly upregulated and downregulated genes (adjusted p-value < 0.05), using all detected genes in the dataset as the background. Additionally, DEGs from clusters 16 and 17 were profiled using GO and KEGG pathway enrichment analyses with enrichGO and enrichKEGG functions with default parameters. Enrichment results were visualized using functions from the enrichplot (1.28.4) package (44).

### Cell differentiation trajectories

To predict the least differentiated cells and the assumed starting point of differentiation in tumor cells, the CytoTRACE2 algorithm (1.1.0) was used with default settings (45). Clusters 16, 17 and 5 with the highest CytoTRACE2 potency scores in the dataset (more than 0.75) were defined as the root clusters for subsequent trajectory analysis. To investigate possible differentiation trajectories and infer the pseudotemporal cell ordering, the Slingshot algorithm (2.16.0) was used (46). The algorithm was used with the existing cluster labels (reducedDim=”umap”) to define the graph structure of the trajectory. The root clusters (start.clus=c(5,16,17)) were selected based on the highest CytoTRACE potency scores, and cellular distances were computed using dist.method = “mnn”.

### Transcription factor activity inference

Regulatory programs at the transcription factor (TF) level were revealed with the decoupleR package (2.14.0) (47). ms at the transcription factor (TF) level were revealed with the decoupleR package (2.14.0) (48). A reference dataset of the human TF-target gene regulatory network was retrieved with the decoupleR::get_collectri function, containing interactions derived from literature curation and ChIP-seq data. Activity scores were estimated using the univariate linear model method (run_ulm, minsize = 5), as suggested in the vignette. Per-cluster mean TF activities were computed by averaging scaled scores within each Seurat cluster. The top 50 most variable TFs across clusters were identified by ranking the standard deviation of cluster-wise mean activities. Heatmaps of cluster-level TF activities were generated using pheatmap (1.0.13) (48) with column-wise (TF-wise) z-scoring.

### hdWGCNA co-expression network analysis

Single-cell high-dimensional weighted gene co-expression network analysis (hdWGCNA) was performed using the R package hdWGCNA (0.4.07) to examine co-expression modules in cluster 16 of the integrated tumor cells dataset (49, 50). A signed co-expression network was constructed with a soft-thresholding power (β = 18), which was selected based on approximate scale-free topology (R² > 0.8) using TestSoftPowers. The hdWGCNA analysis was performed following the default workflow, including network construction (ConstructNetwork), followed by module detection via hierarchical clustering and dynamic tree cutting (default parameters). Module eigengenes (MEs) and harmonized module eigengenes (hMEs) were computed across all cells using ModuleEigengenes with sample-level harmonization. Intramodular connectivity (kME) was calculated within the target population using ModuleConnectivity. Hub genes were identified as the top 25 genes ranked by ModuleConnectivity within each module. Module activity scores were computed using UCell on the top 25 hub genes per module (ModuleExprScore, method = “UCell”). Differential module activity across slingshot pseudotime was evaluated using the built-in PlotModuleTrajectory function.

The co-expression modules were projected onto the Visium datasets using the ProjectModules function with default parameters. Spatial visualizations were generated using SpatialFeaturePlot based on values from the GetMEs function, which returns the connectivity values for the co-expression module.

### Publicly available spatial transcriptomic datasets

The processed Visium FF lung adenocarcinoma No.3 sample was retrieved from the DBKERO database (51). Filtered matrices and tissue images of healthy patients with no known lung disease were retrieved from the BioStudies portal (accession number S-BSST1410) and included samples V10T31-019-A1 (non-smoker male), V10T31-015-A1 (ex-smoker male), and V19S23-092-A1 (smoker female) (52).

### Spatial transcriptomics data processing and annotation transfer

The spatial datasets were imported into the R programming environment (4.5.1) for further analysis with Seurat (5.3.0). The FF lung adenocarcinoma No. 3 sample from the DBKERO database was provided as pre-filtered and was used without additional quality control filtering. For samples from BioStudies S-BSST1410, spots with nFeature_Spatial > 250, nCount_Spatial > 500, and percent.mt < 10% were retained for downstream analysis. Normalization was performed using SCTransform. Dimensionality reduction using PCA, clustering, and UMAP visualization were performed using the first 30 PCs.

Reference-based label transfer was performed using Seurat’s anchor-based integration framework. Cluster identities corresponding to clusters 16 and 17 obtained from scRNA datasets were transferred to the spatial datasets using the TransferData function.

### Validation of Module 17 in a pan-cancer sсRNA-seq dataset

To validate the behavior of Module 17 in an independent dataset, a publicly available preprocessed pan-cancer scRNA-seq atlas of 355 941 cells was utilized (53, 54). The atlas was loaded as a fully normalized and annotated Seurat object with precomputed UMAP coordinates, malignant/non-malignant labels, and detailed cell type classification.

Module 17, previously identified in the discovery dataset using the hdWGCNA workflow, was transferred to the pan-cancer atlas using the ProjectModules function. The WGCNA-derived gene weights of Module 17 were mapped onto the external dataset, and the module eigengene was recalculated with GetMEs. The resulting per-cell eigengene values were stored as Modules17 and used as the Module 17 activity score in all downstream analyses.

To identify cells with high Module 17 activity, a quantile-based thresholding strategy was applied. The distribution of Module 17 values was assessed across all cells in the atlas, and the 95th percentile was selected as the cutoff defining the Module 17-high population.

Conserved markers distinguishing malignant and non-malignant Module 17-high cells were identified using the FindConservedMarkers function, with cancer type set as the grouping variable. For each cancer type, differential expression was calculated using logfc.threshold = 0.25 and min.pct = 0.1. Genes were retained only if adjusted p-values were < 0.05 across all cancer types, and if the detection ratio (pct.1/pct.2) was ≥ 2 consistently in each cancer type.

### Statistics

Data visualization was performed using the ggplot2 package (3.5.2) (55). The distribution of CytoTRACE2 scores across clusters was illustrated using a bar plot, where the height of each bar represents the mean CytoTRACE2 score for each cluster. Error bars were added to represent the standard error of the mean, calculated as the standard deviation divided by the square root of the sample size. Statistical analysis was conducted using the R programming environment (4.5.1). The normality of CytoTRACE2 score distribution was assessed using the Anderson-Darling test, and since it did not follow normal distribution, nonparametric tests were employed. The Kruskal-Wallis rank sum test was applied to compare CytoTRACE2 scores across the clusters, followed by pairwise Wilcoxon rank-sum tests with Benjamini-Hochberg adjustment for multiple testing. Statistical significance was defined as follows: not significant (ns) for p > 0.05, significant at *p < 0.05, **p < 0.01, ***p < 0.001. The enrichment of Module 17-high cells within tumor and normal cells was assessed by constructing a 2×2 contingency table based on atlas annotations, and statistical significance was evaluated using Fisher’s exact test.

## Results

### 1. Characterization of embryoid body and tumor single-cell datasets

To identify common molecular programs shared between embryogenesis and oncogenesis, the analysis included samples representing two distinct biological contexts: normal early human development and malignant growth. As a model of early human embryogenesis, embryoid bodies were generated through spontaneous differentiation of three independent iPSC lines: ICGi022-A (sample EB1), iPS-6637/21 (sample EB2), and iPS-6638/21 (sample EB3). The presence of derivatives from all three germ layers (ectoderm, mesoderm, and endoderm) was confirmed by immunocytochemical staining (Figure 1A-C).

**Figure 1.**
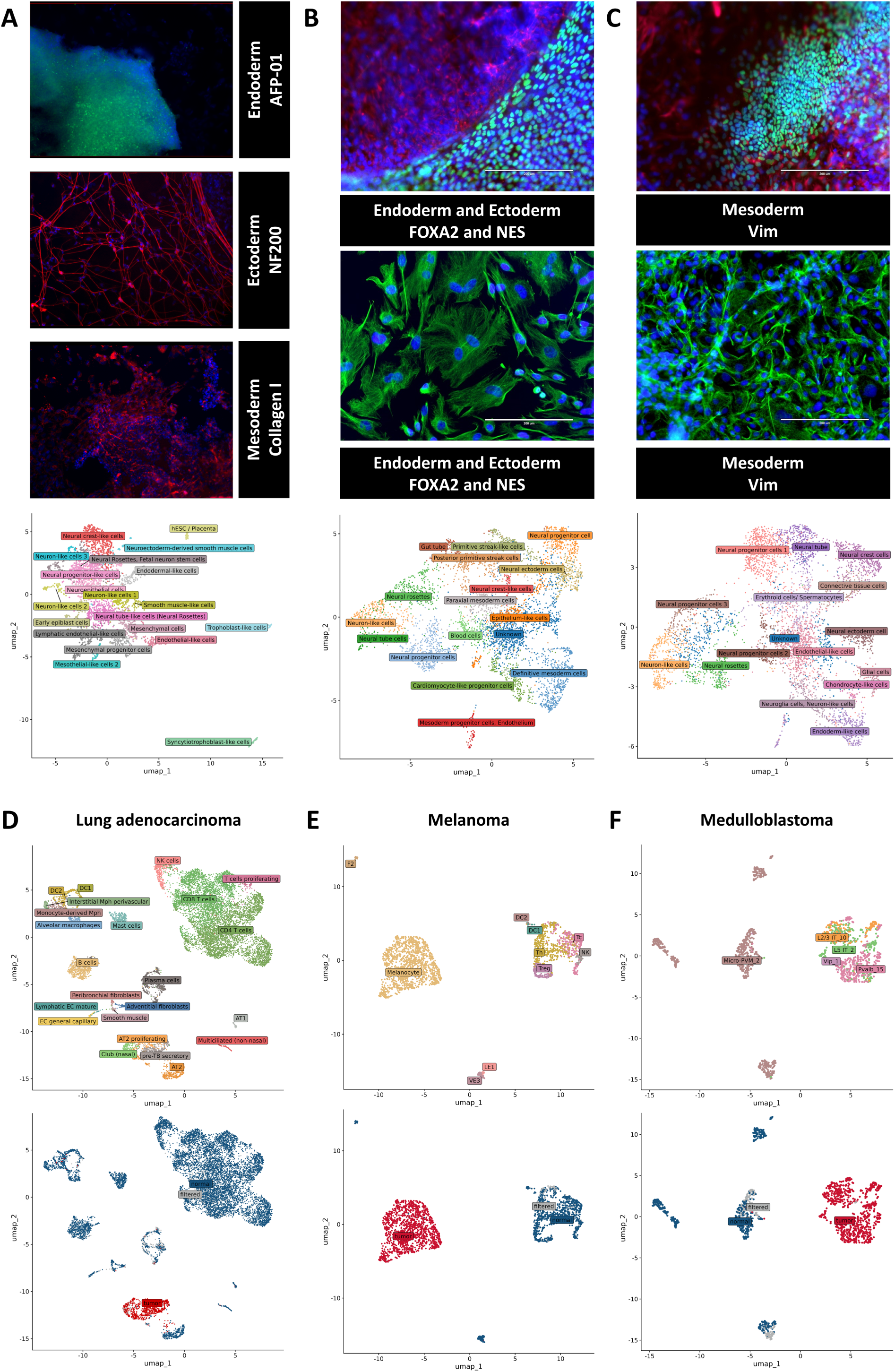
Characterization of tumor and EBs scRNA-seq datasets. Upper panel: Immunofluorescence staining of derivatives representing the three germ layers - ectoderm, mesoderm, and endoderm - in embryoid bodies generated from three independent iPSC lines: ICGi022-A (A), iPS-6637/21 (B), and iPS-6638/21 (C). Middle panel: UMAP visualization of manually annotated cell types across the three embryoid body samples. Lower panel: Representative examples of tumor cell annotation and tumor cell identification for three ontogenetically distinct cancer types. For each cancer type - lung adenocarcinoma (D), melanoma (E), and medulloblastoma (F) - the upper panels show cell type annotations generated using CellTypist, while the lower panels display tumor cell calling based on inferred large-scale copy-number alterations.

Single-cell RNA-seq profiling of all EBs provided a high-resolution view of their cellular architecture, revealing a spectrum of transcriptionally distinct populations corresponding to all three germ layers. Across samples, the EBs contained neuroectoderm-associated populations (including neural progenitors, neural rosettes, neuron-like and neuroepithelial-like cells), mesoderm-associated populations (such as mesenchymal and endothelial-like cells), endoderm-like populations, and multiple intermediate progenitor populations (Figure 1A-C).

To identify molecular programs that are shared between embryogenesis and oncogenesis, the analysis included three ontogenetically and clinically distinct cancer types: lung adenocarcinoma, melanoma, and medulloblastoma. Each tumor dataset was processed independently to identify major cell populations. Initial cell-type annotation was performed using the CellTypist classifier with reference models appropriate for the corresponding cancer type. Tumor cells were subsequently identified by inferring large-scale copy-number alterations, which facilitated the distinction between tumor and normal cells. Representative examples of tumor cell identification for each cancer type are shown in Figure 1D-F.

### 2. Integration of embryoid bodies with different cancers reveals shared cell populations

To determine whether tumors of different tissue origins contain cell populations transcriptionally related to early developmental populations, we performed pairwise integration of EB-derived cells with each cancer type independently, including melanoma, lung adenocarcinoma, and medulloblastoma. Shared clusters were defined as those containing cells from all three EBs samples together with tumor cells from all samples of the corresponding cancer type. This approach allowed us to identify tumor cell populations that converged with EB-derived transcriptional states within each cancer type.

For the melanoma-EBs integration, nine clusters containing cells from all three EBs and all three melanoma samples were identified as shared: 0, 1, 2, 3, 4, 9, 12, 13, and 14 (Figure 2A). Conserved markers were examined first using the combined melanoma and EB-derived cells, filtered with the following thresholds: adjusted p-value < 0.05, log2FC > 0.5, and pct.1 > 0.25. This analysis highlighted distinct signatures: elevated expression of *LGALS3* in cluster 4, DNA replication and repair-associated genes (*TYMS*, *PCLAF*, *DUT*, *PCNA*, *HIST1H4C*) in cluster 12, and mitotic regulators (*ASPM*, *CDC20*, *UBE2C*, *CCNB1*, *MAD2L1*) in cluster 14 (Supplementary Table 3).

**Figure 2.**
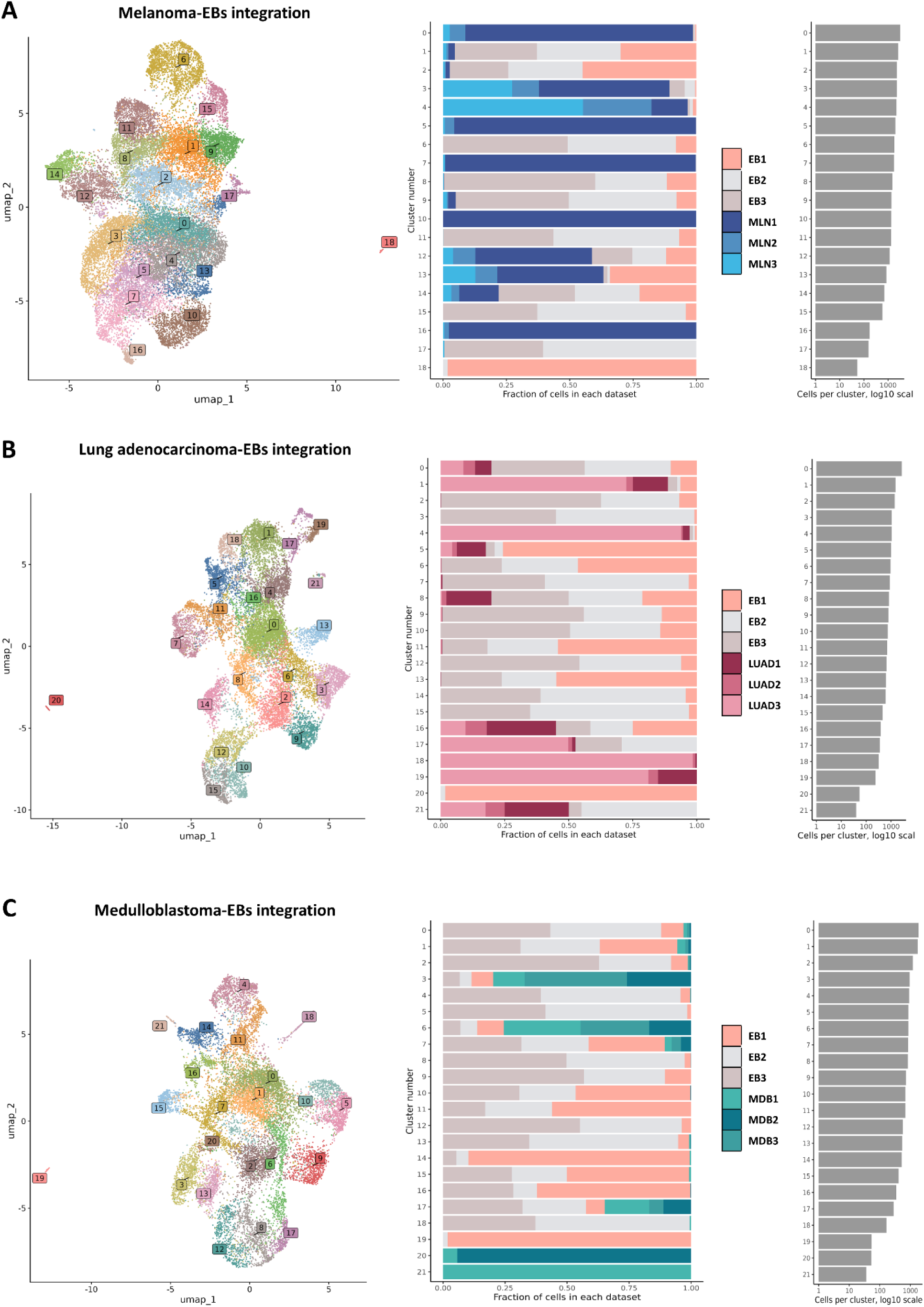
Integration of embryoid body-derived cells with tumor cells from three distinct cancer types. For each cancer type - melanoma (A), lung adenocarcinoma (B), and medulloblastoma (C) - three visualizations are shown. Left panel: UMAP embeddings of the integrated tumor-EBs datasets with cluster labels. Middle panel: fraction of cells contributed by each sample to each cluster. Right panel: total number of cells per cluster on a log10 scale.

To further resolve melanoma-specific conserved programs within these shared clusters, differential expression analysis was repeated using only melanoma-derived cells (adjusted p-value < 0.05, log2FC > 0.5, pct.1 > 0.5). This revealed transcriptional regulators (*ID2*, *RGS2*, *HES1*) in cluster 3, translation-associated genes (*EEF1G*, *RPS3A*, *RPL5*, *RPS4X*) in cluster 4, DNA replication factors (*RRM2*, *CDC45*, *TYMS*, *TK1*, *PCLAF*) in cluster 12, interferon-stimulated *IFIT* genes in cluster 13, and mitotic regulators (*PLK1*, *NEK2*, *DLGAP5*, *UBE2C*, *MKI67*) in cluster 14 (Supplementary Table 4).

For the lung adenocarcinoma-EBs integration, clusters 1 and 8 contained cells from all three EBs and all three cancer samples (Figure 2B). Conserved markers were first evaluated using the combined tumor-and EB-derived cells (adjusted p-value < 0.05, log2FC > 0.5, pct.1 > 0.25). Cluster 1 showed epithelial and adhesion-associated genes (*GRHL2*, *SVIL*), whereas cluster 8 displayed a proliferative program including *BIRC5*, *UBE2C*, *TOP2A*, *NUSAP1*, and *CDK1* (Supplementary Table 5).

To identify markers of lung adenocarcinoma-derived cells within these clusters, analysis was repeated using only tumor-derived cells (adjusted p-value < 0.05, log2FC > 0.5, pct.1 > 0.5). This revealed dominant expression of ribosomal genes in cluster 0, regulators (*NR3C2*, *FTX*, *ARHGAP24*, *SAMD4A*) in cluster 1, the cytoskeletal gene *FLNA* in cluster 5, and core proliferative genes (*UBE2C*, *TOP2A*, *TK1*, *ZWINT*) in cluster 8 (Supplementary Table 6).

For the medulloblastoma-EBs integration, clusters 3, 7, and 17 contained cells from all three EBs and all three cancer samples (Figure 2C). Conserved markers were first evaluated using the combined tumor-and EB-derived cells (adjusted p-value < 0.05, log2FC > 0.5, pct.1 > 0.25). In this analysis, cluster 3 expressed neuronal structural genes (*STMN2*, *TUBB2B*, *MLLT11*, *TUBB3*, *TUBA1A*), cluster 7 displayed mitotic and proliferative regulators (*DLGAP5*, *CENPE*, *BIRC5*, *ASPM*, *TOP2A*), and cluster 17 was marked by the synaptic adhesion gene *NRXN1* (Supplementary Table 7).

To identify transcriptional programs in medulloblastoma-derived cells, differential expression analysis was repeated using only tumor-derived cells (adjusted p-value < 0.05, log2FC > 0.5, pct.1 > 0.5). This revealed DNA replication and repair genes (*EXO1*, *UHRF1*, *CDC45*, *BRIP1*) in cluster 2, cytoskeletal and chromatin regulators (*MARCKSL1*, *TUBA1A*, *H3F3A*) in cluster 3, neuronal differentiation markers (*MYT1L*, *DACH1*, *TRIO*, *EBF1*, *DSCAML1*) in cluster 6, mitotic genes (*CDC25C*, *KIF18B*, *DLGAP5*, *KIF4A*, *ASPM*) in cluster 7, and synaptic and signaling genes (*CACNA1E*, *GABRB1*, *SGIP1*, *ELMO1*) in cluster 17 (Supplementary Table 8).

Despite the distinct tissue origins of melanoma, lung adenocarcinoma, and medulloblastoma, each cancer type contained tumor cell populations that converged with EBs transcriptional states. Across these independently integrated cancer types, one of the most consistent shared features was a proliferative program marked by the expression of *UBE2C*, *TOP2A*, *BIRC5*, and *NUSAP1* genes.

### 3. Integration of distinct cancer types reveals conserved tumor cell populations transcriptionally similar to embryoid body-derived cells

To determine whether the EB-associated tumor cell populations identified independently in melanoma, lung adenocarcinoma, and medulloblastoma correspond to common populations across these cancers, tumor cells from all three cancer types were jointly integrated. This analysis resolved 21 transcriptional clusters (Figure 3A), several of which contained cells from more than one cancer type.

**Figure 3.**
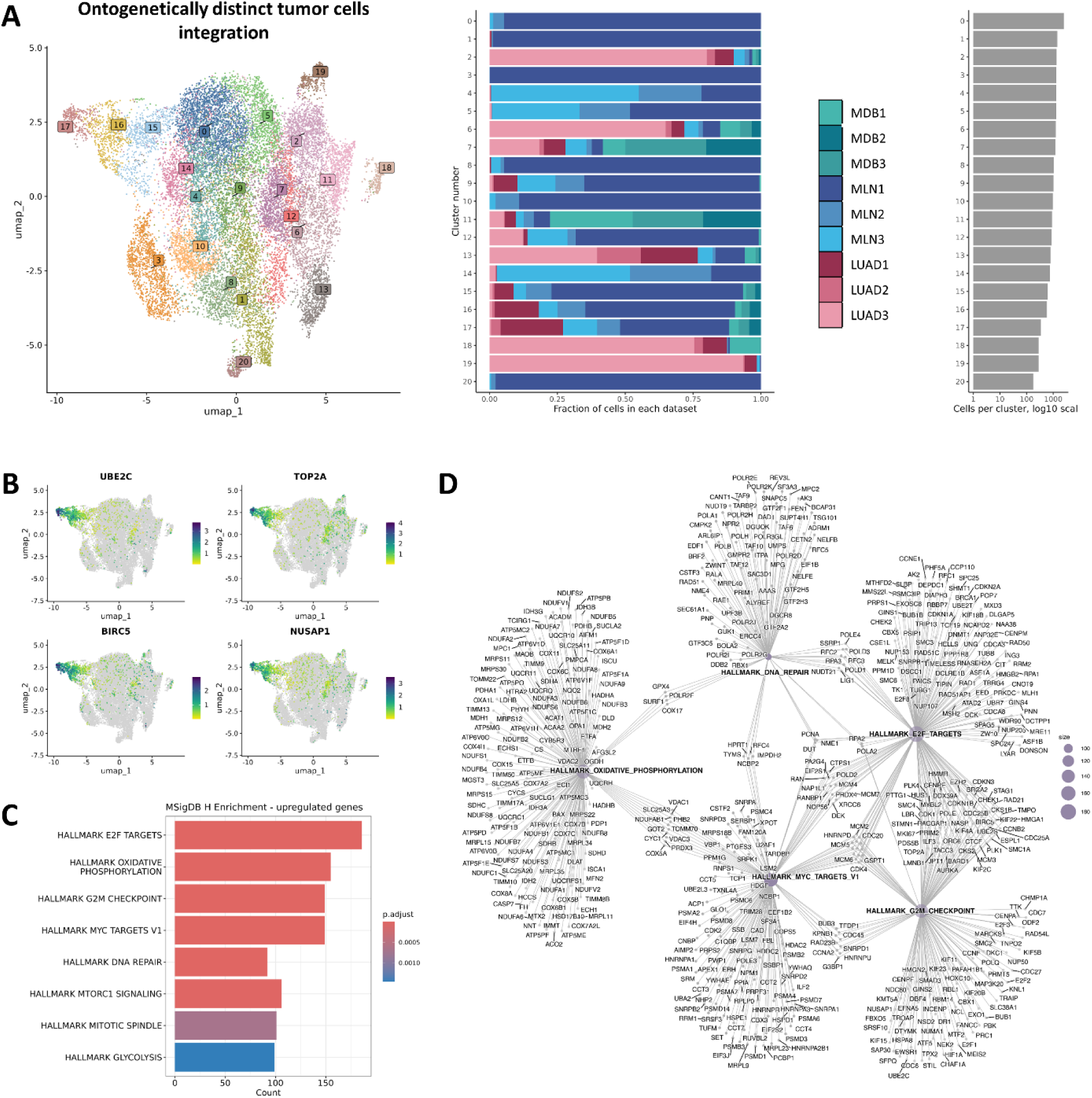
Conserved tumor cell populations shared across lung adenocarcinoma, melanoma, and medulloblastoma. (A) Left: UMAP embedding of tumor cells from lung adenocarcinoma, melanoma, and medulloblastoma following joint integration, resulting in 21 transcriptional clusters. Middle: fraction of cells contributed by each tumor type to every cluster. Right: total number of cells per cluster (log10 scale). (B) Feature plots showing expression of conserved proliferative markers identified in the EBs-tumor integrations (*UBE2C*, *TOP2A*, *BIRC5*, and *NUSAP1*). (C) Hallmark pathway enrichment analysis of genes jointly upregulated in clusters 16 and 17 relative to all other tumor cells. (D) Network representation of enriched Hallmark categories jointly upregulated in clusters 16 and 17, with edges reflecting shared genes between pathways.

The conserved genes identified across the independent tumor-EBs integrations were then examined in the joint tumor atlas. Expression of *UBE2C*, *TOP2A*, *BIRC5*, and *NUSAP1* was concentrated in clusters 16 and 17, where the highest signal was observed (Figure 3B). Both clusters contained tumor cells from melanoma, lung adenocarcinoma, and medulloblastoma, indicating that these tumor cell populations were shared across cancers of distinct tissue origins rather than restricted to a single tumor lineage (Supplementary Figure 1A). We therefore refer to the populations represented by clusters 16 and 17 as conserved tumor cell populations shared across ontogenetically distinct cancers.

To assess whether clusters 16 and 17 were driven primarily by cell cycle-associated variation, an additional control integration was performed with cell cycle-associated variation regressed out. Under these conditions, clusters corresponding to the original clusters 16 and 17 remained distinct and clearly separated in the UMAP, rather than dispersing across the embedding (Supplementary Figure 1B-E). This observation suggests that clusters 16 and 17 are not merely a consequence of cell cycle activity, but represent distinct conserved tumor cell populations across distinct tumor types.

### 4. Functional characterization of the conserved tumor cell populations

To characterize the functional programs associated with the conserved tumor cell populations, enrichment analyses were performed separately for clusters 16 and 17. Cluster 16 demonstrated strong enrichment for cell-cycle-associated GO categories, including chromosome segregation, nuclear and mitotic division, organelle fission, and DNA replication (Supplementary Figure 2A). KEGG pathway analysis was consistent with this proliferative profile, with cell cycle emerging as the top pathway and additional hits corresponding to replication-associated and DNA damage response pathways (Supplementary Figure 2B).

Cluster 17 showed a similarly pronounced proliferative profile. GO analysis highlighted chromosome segregation, mitotic nuclear division, sister chromatid segregation, organelle fission, and microtubule cytoskeleton organization involved in mitosis (Supplementary Figure 2C). KEGG analysis identified the cell cycle as the dominant pathway, together with enrichment of G2/M checkpoint, p53 signaling, and cellular senescence pathways (Supplementary Figure 2D).

To further characterize the shared transcriptional program of clusters 16 and 17, we performed Hallmark enrichment analysis using the genes jointly upregulated in these clusters relative to all other tumor cells (Figure 3C). The combined DEG set showed significant enrichment for HALLMARK_E2F_TARGETS, HALLMARK_G2M_CHECKPOINT, HALLMARK_MYC_TARGETS_V1, HALLMARK_DNA_REPAIR, HALLMARK_MITOTIC_SPINDLE, HALLMARK_OXIDATIVE_PHOSPHORYLATION, and HALLMARK_MTORC1_SIGNALING. To assess how these pathways relate to one another, we constructed a network representation linking enriched Hallmark categories through their shared genes (Figure 3D). The network revealed that HALLMARK_DNA_REPAIR and HALLMARK_MYC_TARGETS_V1 formed major connectivity hubs, each displaying links to all other enriched Hallmark programs. In contrast, HALLMARK_E2F_TARGETS formed a strong but more selectively connected node, linked primarily to HALLMARK_G2M_CHECKPOINT and HALLMARK_MYC_TARGETS_V1, but not connected to HALLMARK_OXIDATIVE_PHOSPHORYLATION.

Together, these analyses indicate that the conserved tumor cell populations represented by clusters 16 and 17 are characterized by coordinated activation of cell cycle, DNA replication, and mitosis-associated transcriptional programs across melanoma, lung adenocarcinoma, and medulloblastoma.

### 5. E2F-TFDP transcription factor axis in the conserved tumor cell populations

To identify upstream regulators of the conserved tumor cell populations, we inferred transcription-factor activity using the decoupleR framework and ranked TFs by normalized enrichment score (NES). The activity heatmap of the top 50 most variable transcription factors revealed a clear segregation of clusters 16 and 17 from all other tumor cell populations (Figure 4A). This separation was associated with elevated inferred activity of cell-cycle-associated TFs, most prominently E2F1, E2F4, and their dimerization partner TFDP1, which showed the highest activity levels (NES > 1.5, p < 0.05) (Figure 4B-D). Elevated inferred activity of additional proliferation-related regulators, including MYC, FOXM1, PLAGL2, and HINFP, further supported the presence of an E2F-centered transcriptional program.

**Figure 4.**
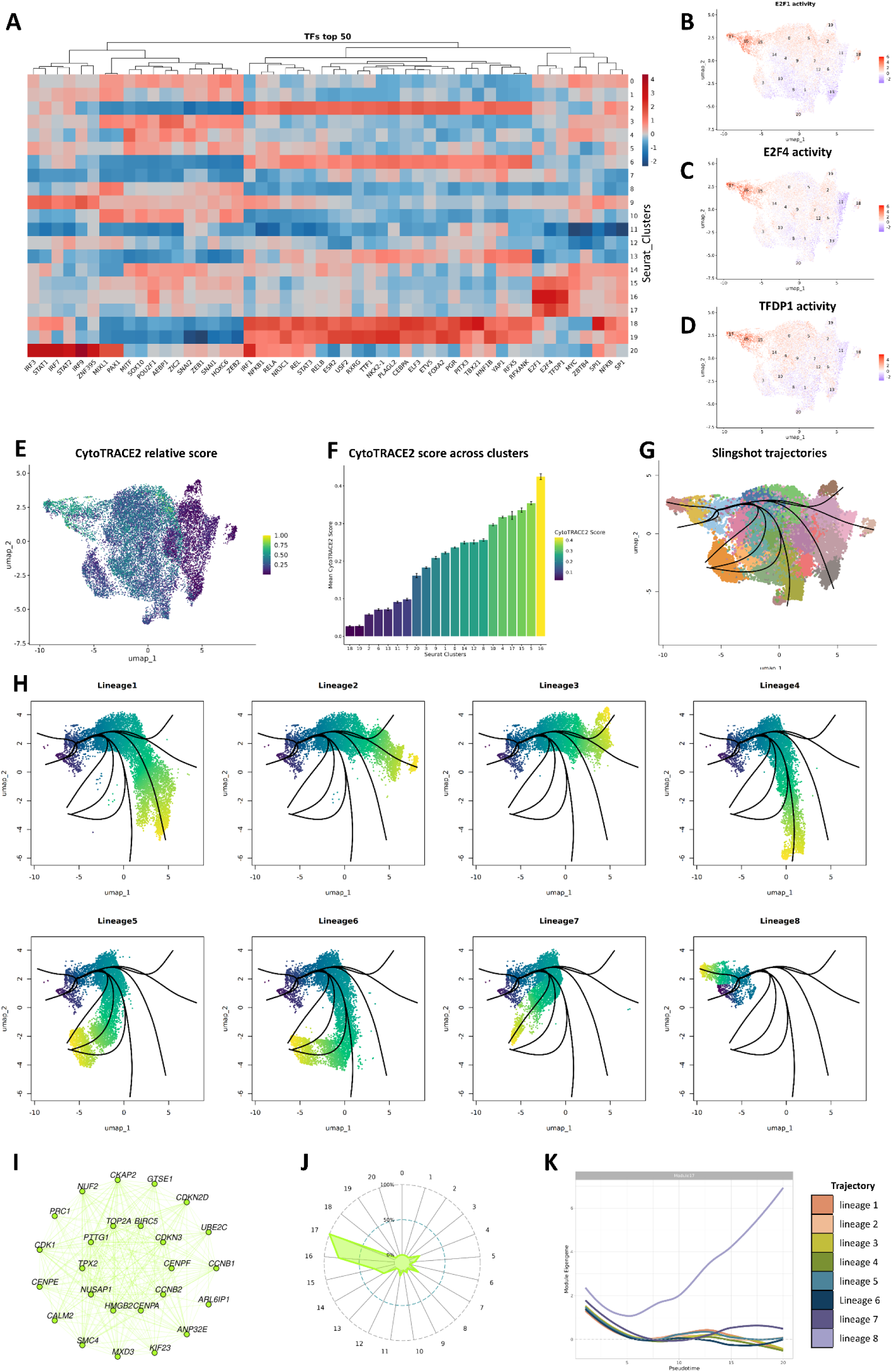
E2F-TFDP-associated conserved tumor cell populations, inferred developmental hierarchy, and transcriptional characteristics of co-expression Module 17. (A) Heatmap of transcription-factor activity scores (top 50 most variable TFs) inferred using decoupleR. (B-D) UMAP projections of inferred activity for E2F1, E2F4, and TFDP1. (E) UMAP visualization of CytoTRACE2 scores across all cells. (F) Distribution of CytoTRACE2 scores across clusters, shown as mean values with standard error bars. (G) Slingshot-inferred global lineage structure with cluster 16 identified as the designated root cluster. (H) Eight pseudotime lineages reconstructed by Slingshot. (I) Network representation of the top 25 hub genes. (J) Module 17 activity across tumor cell clusters (UCell scoring). (K) Changes in Module 17 activity along pseudotime.

In contrast, TFs associated with immune signaling (e.g., IRF, STAT) and differentiation-related factors (e.g., MITF, CEBPA, HOXC6) showed reduced inferred activity in clusters 16 and 17. This transcription factor activity profile is consistent with a poorly differentiated state characterized by enrichment of proliferative transcriptional programs and reduced inferred activity of lineage-associated factors.

Together, these analyses show that the conserved tumor cell populations represented by clusters 16 and 17 are characterized by a distinct inferred transcription factor activity profile, with prominent E2F1, E2F4, and TFDP1 activity accompanied by reduced activity of lineage-associated regulators.

### 6. Pseudotime trajectory inference identifies cluster 16 as an early conserved tumor cell population

To reconstruct putative differentiation hierarchies within the tumor cell compartment, we first applied CytoTRACE2 to rank clusters based on their inferred developmental potential. Clusters 5, 16, and 17 exhibited the highest CytoTRACE2 scores (> 0.75), consistent with lower differentiation states (Figure 4E). A Kruskal-Wallis test confirmed significant differences in CytoTRACE2 scores across all clusters (p < 2.2 × 10⁻¹⁶). Pairwise comparisons further identified clusters 5, 16, and 17 as having significantly higher CytoTRACE2 scores compared to other clusters, consistent with a poorly differentiated phenotype. These clusters exhibited the highest mean CytoTRACE2 scores, supporting their characterization as less differentiated tumor cell populations with high inferred developmental potential (Figure 4F, Supplementary Figure 3A,B). Next, we performed trajectory inference using Slingshot on the RPCA-integrated UMAP embedding, considering clusters 5, 16, and 17 as potential lineage origins. These clusters were chosen because they exhibited the highest CytoTRACE2 scores and therefore were inferred to represent the least differentiated populations in the integrated tumor dataset. Slingshot inferred cluster 16 as the global root, reconstructing eight major pseudotime lineages that radiated outward along the UMAP manifold (Figure 4G). These lineages exhibited smooth trajectories spanning the transcriptional states observed in the tumor compartment, from less differentiated to more differentiated states.

Examination of the individual trajectories further clarified the relative positions of clusters 16 and 17 (Figure 4H, Table 1). Across these trajectories, cluster 16 occupied the earliest pseudotime position, supporting its placement among the least differentiated tumor cell populations identified in the integrated dataset. Notably, cluster 17, despite its high CytoTRACE2 score, was positioned immediately downstream of cluster 16 in lineage 8, suggesting that it may represent a proliferative intermediate within the inferred differentiation trajectories.

**Table 1.** Slingshot-inferred pseudotime lineages across tumor cell populations.

| Slingshot lineage | Pseudotime cluster path |
|---|---|
| 1 | 16 → 15 → 0 → 5 → 12 → 7 → 6 → 13 |
| 2 | 16 → 15 → 0 → 5 → 2 → 11 → 18 |
| 3 | 16 → 15 → 0 → 5 → 2 → 19 |
| 4 | 16 → 15 → 0 → 9 → 1 → 20 |
| 5 | 16 → 15 → 0 → 4 → 10 → 3 |
| 6 | 16 → 15 → 0 → 9 → 1 → 8 |
| 7 | 16 → 15 → 0 → 14 |
| 8 | 16 → 17 |

### 7. hdWGCNA reveals proliferation-associated module in cluster 16

To characterize coordinated transcriptional programs within the conserved tumor cell population of interest with high inferred developmental potential, we performed high-dimensional weighted gene co-expression network analysis on cluster 16 from the integrated multi-cancer scRNA-seq atlas. A signed co-expression network was constructed, resulting in 25 distinct co-expression modules (Supplementary Figure 4). Among these, Module 17 emerged as the most biologically relevant due to its strong association with known proliferation markers. This module contained established proliferation markers such as *TOP2A*, *MKI67*, *BIRC5*, *UBE2C*, and *CENPF*.

To explore the gene network further, we examined the top 25 hub genes in this module, which were highly interconnected and included key regulators of mitosis, such as *TOP2A*, *BIRC5*, *PRC1*, *CDK1*, *CENPF*, *CENPE*, *SMC4*, and *KIF23*, all of which are crucial for mitotic spindle function, chromosome alignment, segregation, and overall cell cycle progression (Figure 4I). These genes were subsequently defined as the 25-gene proliferation-associated signature. We further explored the functional significance of this gene set and found that it was strongly enriched for biological pathways related to Mitotic Spindle, G2/M checkpoint, and E2F targets, as defined by the Hallmark MSigDB 2020 (Supplementary Figure 2E). To confirm the module’s association with a proliferative state, we performed UCell-based scoring of the 25 hub genes. This analysis revealed strong module activity in cluster 16 and, to a lesser extent, in cluster 17 (Figure 4J). Consistent with these findings, a UMAP projection of Module 17 activity further showed that its signal was concentrated almost exclusively within clusters 16 and 17, reinforcing the conclusion that Module 17 activity is largely restricted to the conserved tumor cell populations (Supplementary Figure 5).

In addition, we evaluated the expression of this 25-gene proliferation-associated signature in the context of pseudotime analysis. Trajectory analysis using Slingshot, initiated from clusters with high CytoTRACE2 scores (including cluster 16), revealed that the activity of Module 17 peaked early in pseudotime and progressively declined along differentiation lineages (Figure 4K). This pattern is consistent with a proliferation-associated, progenitor-like transcriptional profile at early pseudotime positions, suggesting that the signature marks tumor cell populations with high proliferative activity and inferred developmental potential.

### 8. Spatial validation of conserved tumor cell populations and their 25-gene proliferation-associated signature

To explore the spatial organization and microenvironmental context of the conserved tumor cell populations represented by clusters 16 and 17, we transferred cluster labels onto Visium spatial transcriptomics datasets from lung adenocarcinoma samples and lung tissue from individuals without known lung disease (non-smoker male, ex-smoker male, and smoker female).

In the lung adenocarcinoma No.3 Visium sample, anchor-based label transfer (FindTransferAnchors followed by TransferData) revealed a consistent, spatially restricted distribution of predicted cluster 16 and 17 signals. Prediction scores indicated preferential localization of these populations within rosette-like epithelial structures with expanded luminal spaces (Figure 5A). Histologically, these structures were characterized by enlarged lumina, often filled with homogeneous eosinophilic material resembling edematous fluid, sometimes mixed with fresh erythrocytes. The tumor cells in these regions appeared morphologically similar to adjacent cells but were slightly smaller with occasional hyperchromatic nuclei. The morphological distinction was more prominent than the cytological one: adjacent tumor cells typically formed solid nests or classical rosettes, while regions with elevated prediction scores for clusters 16 and 17 displayed rosettes with wider luminal spaces and looser epithelial arrangements along their margins.

**Figure 5.**
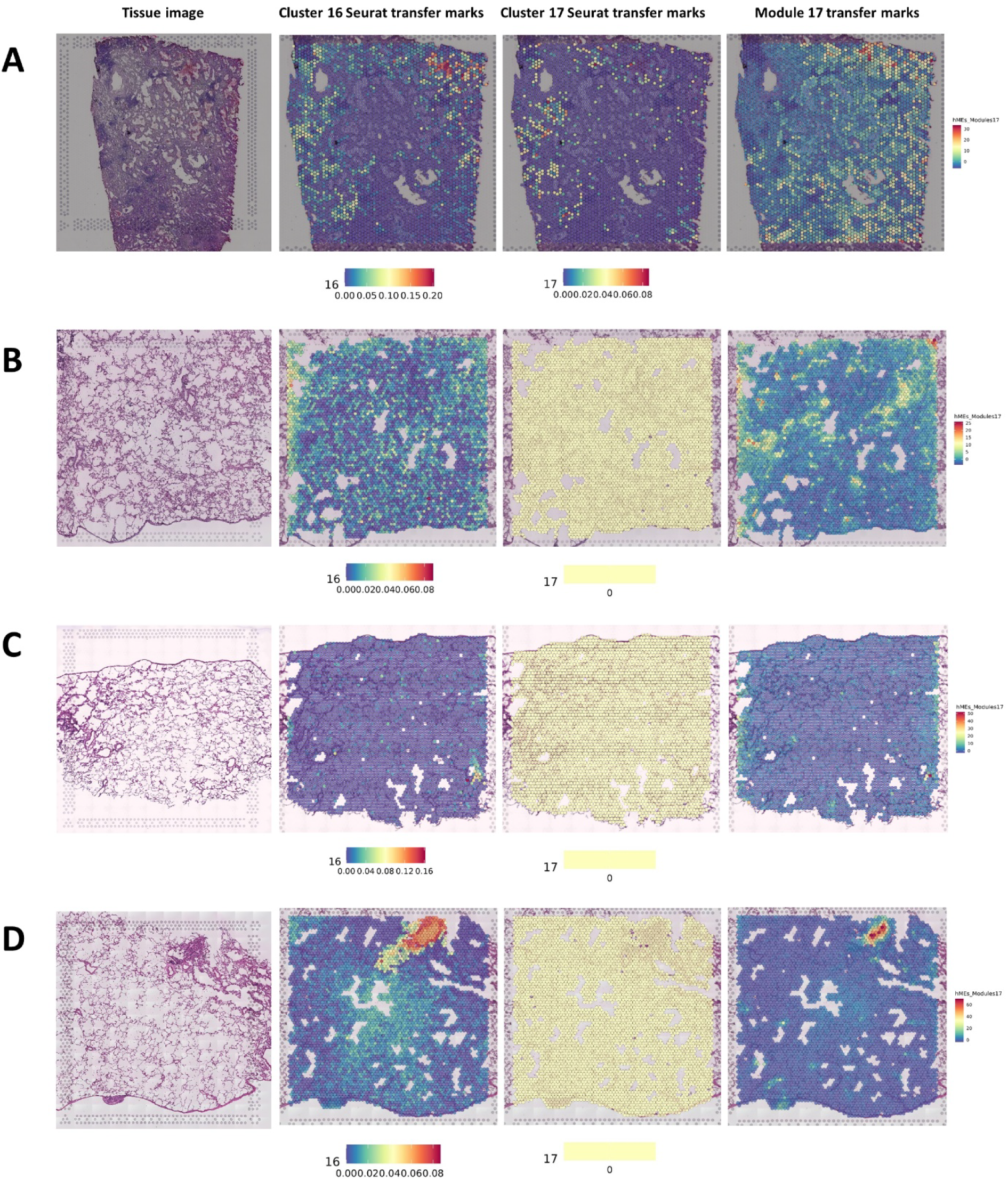
Spatial distribution of conserved tumor cell populations and the 25-gene proliferation-associated signature in lung adenocarcinoma and non-diseased lung samples. A-D: Tissue images of lung samples (first column) from different conditions: (A) Lung adenocarcinoma No.3 tissue image, (B) Non-diseased lung tissue from a non-smoker male, (C) Non-diseased lung tissue from an ex-smoker male, (D) Non-diseased lung tissue from a smoker female. The second, third, and fourth columns display the spatial distributions of prediction scores for cluster 16, cluster 17, and the 25-gene proliferation-associated signature, respectively.

Although these structures are not typically considered distinct morphological subtypes in diagnostic practice, their spatial association with predicted conserved tumor cell populations and the proliferation-associated signature suggests that they may correspond to a transcriptionally distinct population within the tumor cell compartment. This finding, combined with the observed morphological differences from surrounding tumor tissue, suggests that these regions warrant further investigation.

Additionally, data from the lung adenocarcinoma No. 3 sample were integrated with the hdWGCNA results to visualize the spatial distribution of the proliferative Module 17. We projected the previously identified co-expression modules onto the Visium dataset, allowing us to visualize the spatial localization of Module 17 activity. The spatial projection revealed that Module 17 activity was elevated near the regions with high cluster 16 transfer prediction scores. This spatial distribution reflected the restricted enrichment of the proliferation-associated program in specific tumor regions, supporting the hypothesis that Module 17 is associated with cluster 16 (Figure 5A). Taken together, these findings indicate that Module 17 is intricately linked to the proliferative transcriptional profile of tumor cells. The spatially restricted activity of Module 17 in these tumor regions highlights its potential as a molecular signature of cell division and tumor growth.

In two non-diseased lung Visium datasets (non-smoker male and ex-smoker male), prediction scores for clusters 16 and 17 were either low or undetectable (Figure 5B,C). This suggests that transcriptional profiles resembling these conserved tumor cell populations were not detected in these samples. However, in the smoker female dataset, elevated prediction scores for cluster 16 and the 25-gene proliferation-associated signature were observed in nearby regions (Figure 5D). Importantly, histological assessment of the same tissue revealed basal cell hyperplasia, a reactive expansion of the airway basal cell compartment that can occur in response to chronic epithelial injury.

In summary, spatial transcriptomics localized the conserved tumor cell populations and their 25-gene proliferation-associated signature to discrete regions of lung adenocarcinoma tissue, while cluster 16-associated signals and the signature were also detected in a region of basal cell hyperplasia in the lung of a smoker without known lung disease.

### 9. Validation of the 25-gene proliferation-associated signature in a pan-cancer scRNA-seq dataset

To assess whether the 25-gene proliferation-associated signature derived from Module 17 represents a recurrent feature across a broader spectrum of human cancers, we evaluated this signature in an independent pan-cancer single-cell RNA-seq atlas (Zenodo record 14511579).

The dataset includes 355 941 tumor and normal cells across eight tumor types: melanoma, melanoma brain metastasis, basal cell carcinoma, triple-negative breast cancer, HER2+ breast cancer, ER+ breast cancer, clear-cell renal cell carcinoma, hepatocellular carcinoma, and intrahepatic cholangiocarcinoma. The atlas was provided as a preprocessed Seurat object with cell-type annotations, malignant/non-malignant classification, and UMAP coordinates (Figure 6A, B).

**Figure 6.**
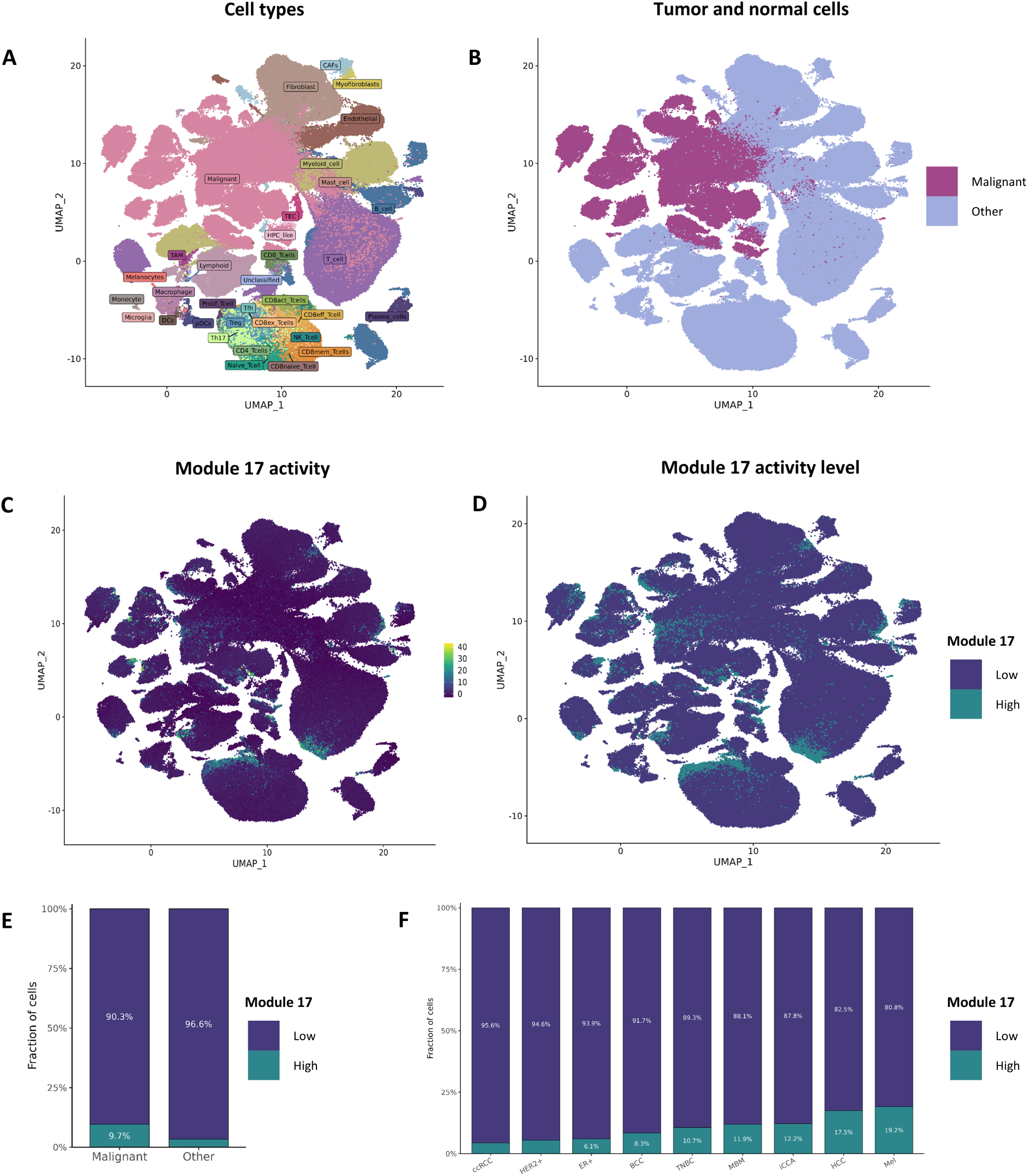
Validation of the activity of the 25-gene proliferation-associated signature derived from Module 17 in a pan-cancer single-cell RNA-seq atlas. (A) Detailed cell type annotation. (B) Tumor and normal cells. (C) 25-gene proliferation-associated signature activity across the pan-cancer atlas. (D) Signature-high and signature-low cells based on a 95th percentile threshold of signature activity. (E) Fraction of signature-high and signature-low cells among tumor and normal cells. (F) Fraction of Module 17-high and Module 17-low cells across cancer entities. ccRCC - clear-cell renal cell carcinoma; HER2+ - HER2+ breast cancer; ER+ - ER+ breast cancer; BCC - basal cell carcinoma; TNBC - triple-negative breast cancer; MBM - melanoma brain metastasis; iCCA - intrahepatic cholangiocarcinoma; HCC - hepatocellular carcinoma; Mel - melanoma.

Scores for the 25-gene proliferation-associated signature were calculated for each cell in the pan-cancer atlas, and their distribution was used to define high signature activity (Figure 6C). A threshold at the 95th percentile of signature scores was applied, and cells exceeding this cutoff were classified as signature-high. This provided a consistent criterion for identifying cells with the highest scores for the 25-gene proliferation-associated signature.

Signature-high cells were detected among both tumor and normal cells within the pan-cancer atlas (Figure 6D). At the same time, their representation differed between these populations, with a higher fraction of signature-high cells observed among tumor cells compared with normal cells (Figure 6E; Fisher’s exact test: odds ratio = 3.04, 95% CI 2.95-3.13, p < 2.2 × 10⁻¹⁶). Importantly, signature-high cells were present in all cancers included in the atlas, although their relative abundance varied between cancer types (Figure 6F). In addition, signature-high cells were not evenly distributed across cellular populations, being more frequently detected in proliferative cell states and only rarely observed in most differentiated cell types (Supplementary Figure 6).

To identify conserved transcriptional features distinguishing tumor signature-high cells from normal signature-high cells across cancer types, differential expression analysis was performed using the following criteria: adjusted p-value < 0.05, log2FC > 0.25, and a detection ratio (pct.1/pct.2) ≥ 2. This analysis identified 14 genes (*EFNA1*, *ERBB3*, *HES1*, *YAP1*, *MDK*, *CTTN*, *TOM1L1*, *RTKN*, *PFN2*, *TSPAN6*, *S100A13*, *S100A16*, *TUSC3* and *CRNDE*) (Supplementary Table 9).

Taken together, these findings indicate that the 25-gene proliferation-associated signature derived from Module 17 is detectable in both normal and tumor cells across multiple cancer types, indicating that this transcriptional program is not restricted to tumor cells. However, cells with high signature activity are significantly more frequent among tumor cells, and tumor signature-high cells exhibit increased expression of specific genes compared with normal signature-high cells.

## Discussion

Oncogenesis recapitulates several features of embryonic development, including rapid proliferation, cellular plasticity, migration, and tissue remodeling (18, 56, 57). Developmental regulators such as PAX family members, DUX4, Kaiso, and noncanonical Wnt signaling are also implicated in malignant transformation (56–58). However, these parallels have largely been inferred from indirect comparisons rather than from analyses integrating embryonic and tumor transcriptional states within a common framework. Here, we directly compared single-cell transcriptional states in lung adenocarcinoma, melanoma, and medulloblastoma with those observed in EBs. Despite their distinct tissue and developmental origins, all three cancer types contained tumor cell populations that reproducibly showed transcriptional similarity to EB-derived states, supporting the existence of conserved tumor cell states across otherwise unrelated cancers.

A prominent feature of these shared populations is high proliferative activity, accompanied by recurrent expression of the cell-cycle-associated genes *UBE2C, TOP2A, BIRC5,* and *NUSAP1*. However, proliferation alone did not account for their transcriptional identity: the populations remained distinguishable after accounting for cell-cycle-associated variation and were characterized by low differentiation status, high inferred developmental potential and early positioning along inferred differentiation trajectories. Together, these findings indicate that the conserved tumor cell populations combine pronounced proliferative activity with low differentiation status rather than reflecting cell-cycle activity alone.

These conserved populations with high inferred developmental potential are characterized by coordinated activity of E2F1, E2F4, and TFDP1. The E2F-TFDP axis regulates DNA replication and cell-cycle progression and is also involved in the proliferation of stem and progenitor cells. TFDP1 and E2F4 are required for efficient proliferation of human hematopoietic stem and progenitor cells (59), while increased E2F-TFDP activity is associated with cancer stem-like phenotypes, chemoresistance, and expression of cell-cycle genes such as *TOP2A* and *BIRC5* in several cancer types (60–62). Thus, the coordinated E2F-TFDP activity in the conserved populations is consistent with the reported roles of this axis in stem and progenitor cell proliferation and in cancer stem-like phenotypes, although our data do not establish that this transcription-factor activity determines their proliferative and low-differentiation characteristics.

Another feature shared by the conserved populations is a 25-gene proliferation-associated signature that includes several mitotic regulators, such as *BIRC5, HMGB2, CDKN2D, ANP32E, PTTG1, CENPF,* and *NUSAP1*. Notably, 21 of the 25 genes overlapped with an established pan-cancer G2/M cell-cycle program described by Gavish et al (63). This substantial overlap indicates that the proliferation-associated signature itself is not unique to the populations identified here. What distinguishes these populations is the association of this broadly shared proliferation-associated program with transcriptional similarity to EB-derived states and low differentiation across cancers of distinct origins.

Validation in an independent pan-cancer scRNA-seq atlas demonstrated broad recurrence of the 25-gene proliferation-associated signature across all eight cancer types examined. Signature-high cells were detected in both tumor and normal populations, although they were significantly more frequent among tumor cells. Comparative analysis of tumor and normal signature-high cells identified 14 tumor-associated genes, including *EFNA1*, *ERBB3*, *HES1*, *YAP1*, and *MDK*, associated with tumor growth and proliferation (64–68), *CTTN, TOM1L1, RTKN, PFN2, and TSPAN6*, involved in cell motility and invasive behavior (69–73), and *S100A13* and *S100A16*, implicated in angiogenesis (74, 75). These findings indicate that, although the proliferation-associated signature is broadly conserved and not restricted to tumor cells, its expression in tumor cells is accompanied by additional transcriptional features associated with tumor progression.

Spatial transcriptomic analysis showed that the 25-gene proliferation-associated signature is predominantly detected in lung adenocarcinoma and largely absent from the examined normal lung samples, with the exception of one smoker lung sample, where it was localized to a region of basal cell hyperplasia, a lesion that can precede squamous cell carcinoma of the lung (76). Previous work has shown that carcinogen exposure in the airway epithelium can promote competition among basal cells, clonal expansion, accumulation of transitional epithelial states, and reduced luminal differentiation, thereby contributing to the development of preinvasive lesions and subsequent tumor formation (77). In this context, detection of the proliferation-associated signature in basal cell hyperplasia raises the possibility that this transcriptional program may emerge at a premalignant stage. However, the limited number of spatial samples precludes conclusions regarding a premalignant role and warrants validation in larger cohorts including premalignant and tumor tissues.

Several limitations should be considered. EBs reproduce aspects of early lineage differentiation but do not fully recapitulate the spatial organization and morphogenesis of the human embryo. Comparison with fetal tissues and more advanced stem cell-derived models, including gastruloids, blastoids, and synthetic embryo models, will therefore be important for refining the developmental identity of the populations described here (24). In addition, although the discovery analysis included three biologically distinct cancer types and the 25-gene proliferation-associated signature was subsequently detected across an independent pan-cancer atlas, broader patient cohorts will be required to determine the prevalence and tissue dependence of this signature. Finally, the analyses used to infer differentiation status, pseudotime trajectories, and transcription-factor activity are based on transcriptional associations and therefore cannot establish direct lineage relationships or regulatory causality. Experimental perturbation and lineage-tracing studies will be required to validate these relationships.

## Conclusions

The integrative single-cell analysis of embryoid bodies and three ontogenetically distinct cancers identifies conserved tumor cell populations transcriptionally similar to EB-derived cells that combine high proliferative activity, low differentiation status, and high inferred developmental potential. Their 25-gene proliferation-associated signature recurs across multiple cancer types in an independent pan-cancer atlas and is accompanied by increased expression of 14 cancer-related genes in tumor cells compared normal cells. Altogether, these findings provide a foundation for evaluating diagnostic and therapeutic value of the conserved tumor cell populations across cancers of distinct origins.

## List of abbreviations

EMT: Epithelial-mesenchymal transition
EB: Embryoid body
scRNA-seq: Single-cell RNA sequencing
iPSCs: Induced pluripotent stem cells
bFGF: Basic fibroblast growth factor
DMEM: Dulbecco’s Modified Eagle Medium
KoSR: Knockout Serum Replacement
FBS: Fetal bovine serum
SRA: Sequence Read Archive
PCs: Principal components
UMAP: Uniform Manifold Approximation and Projection
CNV: Copy number variation
DEGs: Differentially expressed genes
PCA: Principal component analysis
GO: Gene Ontology
KEGG: Kyoto Encyclopedia of Genes and Genomes
MSigDB: Molecular Signatures Database
GSEA: Gene Set Enrichment Analysis
TF: Transcription factor
hdWGCNA: High-dimensional weighted gene co-expression network analysis
MEs: Module eigengenes
hMEs: Harmonized module eigengenes
NES: Normalized enrichment score

## Declarations

### Ethics approval and consent to participate

Not applicable

### Consent for publication

Not applicable

### Availability of data and materials

The datasets analysed during the current study are available in the BioStudies repository, https://doi.org/10.6019/S-BSST1410; in the Zenodo repository, https://doi.org/10.5281/zenodo.14511579; in the Sequence Read Archive (SRA) repository, https://www.ncbi.nlm.nih.gov/sra (accession numbers SRP347260, SRP274357 and SRP389673); and in the DBKERO repository, https://kero.hgc.jp/Ad-SpatialAnalysis_2024.html.

The datasets generated during the current study are available from the corresponding author on reasonable request.

### Authors’ contributions

A.A.K. contributed to conceptualization, methodology, formal analysis, validation, data curation, visualization, supervision and writing - original draft. V.G.S. contributed to methodology, formal analysis, validation, visualization and writing - original draft. M.E.M. contributed to methodology, investigation and writing - original draft. D.I.Z. contributed to methodology, investigation and writing - original draft. T.N.K., A.A.G., E.E.K., A.A.Ko., E.B.D., M.S.T. and U.A.B. contributed to investigation. M.A.B. contributed to resources. D.M.L. contributed to validation. N.A.S. and E.V.D. contributed to conceptualization, methodology, supervision, project administration and writing - review & editing. All authors read and approved the final manuscript.

## Acknowledgments

The authors are grateful to S.M. Zakian (Dr. Sci. Biol., Prof., RAS, Head of the Laboratory of Developmental Epigenetics, Institute of Cytology and Genetics, SB RAS), as well as to A.A. Malakhova (Ph.D. Biol.) and E.V. Grigor’eva (Ph.D. Biol.) for providing the induced pluripotent stem cell line ICGi022-A for this research.

## Supplementary Information

Additional file 1:

Format:.docx

Title: Supplementary Figures 1-6

Description: Supplementary Figure 1. Cell cycle regression analysis of proliferative clusters. (A) Fraction of different tumor samples in clusters 16 and 17. (B) UMAP visualization following integration with regression of cell cycle–associated variation, colored by newly assigned cluster labels. (C) UMAP embedding after cell cycle regression, colored by the original cluster labels, illustrating preservation of cluster identity. (D) Same UMAP as in (C), with original clusters 16 and 17 specifically highlighted. (E) UMAP after cell cycle regression showing expression levels of proliferative marker genes UBE2C, TOP2A, BIRC5, and NUSAP1. Supplementary Figure 2. Functional enrichment analysis of pan-cancer proliferative cell states. (A) GO biological process enrichment in cluster 16. (B) KEGG pathway enrichment in cluster 16. (C) GO biological process enrichment in cluster 17. (D) KEGG pathway enrichment in cluster 17. (E) Hallmark gene set enrichment analysis (MSigDB 2020) of the Module 17. Supplementary Figure 3. CytoTRACE2 scores across tumor cell clusters. (A) Pairwise Wilcoxon rank-sum test p-values comparing CytoTRACE2 scores between clusters. (B) Bar plot showing CytoTRACE2 scores for each cluster. Supplementary Figure 4. Co-expression modules identified in cluster 16 tumor cells. Supplementary Figure 5. UMAP projection of Module 17 activity scores (UCell scoring). Supplementary Figure 6. Fraction of Module 17-high and Module 17-low cells across cell type identities.

Additional file 2:

Format:.xlsx

Title: Supplementary Table 1. Antibodies used for immunofluorescence staining of Ebs

Description: List of antibodies used for immunofluorescence staining of embryoid bodies (EBs).

Additional file 3:

Format:.xlsx

Title: Supplementary Table 2. Overview of analyzed tumor samples

Description: Summary of analyzed tumor samples included in the study.

Additional file 4:

Format:.xlsx

Title: Supplementary Table 3. Conserved markers in shared clusters of melanoma-EB integration

Description: List of conserved markers identified in shared clusters during melanoma-EB integration.

Additional file 5:

Format:.xlsx

Title: Supplementary Table 4. Melanoma-specific conserved markers in shared clusters of melanoma-EB integration

Description: List of melanoma-specific conserved markers identified in shared clusters during melanoma-EB integration.

Additional file 6:

Format:.xlsx

Title: Supplementary Table 5. Conserved markers in shared clusters of LUAD-EB integration

Description: List of conserved markers identified in shared clusters during LUAD-EB integration.

Additional file 7:

Format:.xlsx

Title: Supplementary Table 6. LUAD-specific conserved markers in shared clusters of LUAD-EB integration

Description: List of LUAD-specific conserved markers identified in shared clusters during LUAD-EB integration.

Additional file 8:

Format:.xlsx

Title: Supplementary Table 7. Conserved markers in shared clusters of medulloblastoma-EB integration

Description: List of conserved markers identified in shared clusters during medulloblastoma-EB integration.

Additional file 9:

Format:.xlsx

Title: Supplementary Table 8. Medulloblastoma-specific conserved markers in shared clusters of medulloblastoma-EB integration

Description: List of medulloblastoma-specific conserved markers identified in shared clusters during medulloblastoma-EB integration.

Additional file 10:

Format:.xlsx

Title: Supplementary Table 9. DEGs distinguishing tumor versus normal cells in the Module 17 high populations

Description: Differentially expressed genes distinguishing tumor versus normal cells in Module 17 high populations.

